# Rapid propagation of Ca^2+^ waves and electrical signals in a liverwort *Marchantia polymorpha*

**DOI:** 10.1101/2023.10.16.562169

**Authors:** Kenshiro Watanabe, Kenji Hashimoto, Kota Hasegawa, Hiroki Shindo, Yushin Tsuruda, Kamila Kupisz, Mateusz Koselski, Piotr Wasko, Kazimierz Trebacz, Kazuyuki Kuchitsu

**Author notes:** These authors contributed equally (Kenshiro Watanabe, Kenji Hashimoto). Corresponding Author: Kazuyuki Kuchitsu Department of Applied Biological Science, Tokyo University of Science, Noda, Chiba 278-8510 Japan.

## Abstract

In response to both biotic and abiotic stresses, vascular plants transmit long-distance Ca^2+^ and electrical signals from localized stress sites to distant tissues through their vasculature. Various models have been proposed for the mechanisms underlying the long-distance signaling, primarily centered around the presence of vascular bundles. We here demonstrate that the non-vascular liverwort Marchantia polymorpha possesses a mechanism for propagating Ca^2+^ waves and electrical signals in response to wounding. The propagation velocity of these signals was approximately 1-2 mm/s, equivalent to that observed in vascular plants. Both Ca^2+^ waves and electrical signals were inhibited by La^3+^ as well as tetraethylammonium chloride, suggesting crucial importance of both Ca^2+^ channel(s) and K^+^ channel(s) in wound-induced membrane depolarization as well as the subsequent long-distance signal propagation. Simultaneous recordings of Ca^2+^ and electrical signals indicated a tight coupling between the dynamics of these two signaling modalities. Furthermore, molecular genetic studies revealed that a GLUTAMATE RECEPTOR-LIKE (GLR) channel plays a central role in the propagation of both Ca^2+^ waves and electrical signals. Conversely, none of the three two-pore channels (TPCs) were implicated in either signal propagation. These findings shed light on the evolutionary conservation of rapid long-distance Ca^2+^ wave and electrical signal propagation involving GLRs in land plants, even in the absence of vascular tissue.

## Introduction

Despite the absence of a nervous system, plants are equipped with a plethora of sensors to monitor environmental changes and signal transmission mechanisms that span their entire structures. In response to biotic and abiotic stresses, long-distance signals are systemically transmitted from the stressed local sites to distant tissues. Such long-distance signals play important roles in inducing stress responses throughout the body. One type of long-distance signaling has been suggested to involve electrical signals, Ca^2+^ (Gilroy et al. 2016, Hedrich et al. 2016, Johns et al. 2021), reactive oxygen species (ROS) (Miller et al. 2009, Lew et al. 2020), and hydraulic signals (Malone et al. 1991).

Electrical signals are a change in plasma membrane potential induced by the movement of ions into and out of the cell via ion channels/transporters. Various stimuli have been shown to induce electrical signals in Streptophytes including bryophytes and angiosperms (Paszewski et al. 1982, Kupisz et al. 2017, Sibaoka et al. 1969, Scherzer et al. 2019, Hagihara and Toyota 2020). Experiments using ion-selective microelectrodes and calcium-permeable channel inhibitors have indicated that Ca^2+^ influx is an important step in the generation of membrane depolarization (Trȩbacz et al. 1994, Ward et al. 1995, Kikuyama et al. 1997, Vodeneev et al. 2016, Kupisz et al. 2017). In addition to Ca^2+^ influx, Cl^-^ and K^+^ play a role in electrogenesis (Trȩbacz et al. 1994, Trȩbacz et al. 1989, Vodeneev et al. 2016). In addition to electrical signals, given the presence of numerous Ca^2+^ sensor proteins in plants, plasma membrane Ca^2+^ influx into the cytoplasm triggered by a variety of stimuli plays a central role in signal transduction (Kurusu et al. 2013, Kudla et al. 2018, Thor 2019, Hashimoto et al. 2023).

Stimulus-induced membrane depolarization and Ca^2+^ influx into the cytoplasm not only occur in the stimulated cell, but also propagate to distant cells. Propagation of the Ca^2+^ wave has been suggested to involve glutamate-receptor-like (GLR) channels, which are ligand-gated and non-selectively permeable to cations, as well as two-pore channels (TPCs), which are voltage-activated and permeable to cations (Mousavi et al. 2013, Toyota et al. 2018, Nguyen et al. 2018, Choi et al. 2014, Kiep et al. 2015). Although many angiosperm species including *Arabidopsis thaliana* and rice *Oryza sativa* possess sole *TPC1* gene (Furuichi et al. 2001; Kurusu et al. 2004; Peiter et al. 2005), liverworts and mosses possess type 1 (similar to angiosperm TPC1) and type 2 TPC genes (Hashimoto et al. 2022)

In *Arabidopsis thaliana*, electrical signal and Ca^2+^ wave has been shown to propagate at the fastest rate (1-2 mm/s) along the vascular bundles (Mousavi et al. 2013, Toyota et al. 2018, Nguyen et al. 2018, Bellandi et al. 2022). A recent model has put forth the idea that substances diffuse upwards through the vascular bundle from the site of stimulation, leading to alterations in plasma membrane potential and an increase in [Ca^2+^]_cyt_ (Bellandi et al. 2022, Gao et al. 2023).

Signal propagation have also recently been observed in non-vascular bryophytes. In the moss *Physcomitrium patens,* osmotic pressure and H_2_O_2_ evoke Ca^2+^ waves at a much slower propagation rate, approximately 5 µm/s, which is notably slower compared to vascular plants (Storti et al. 2018, Koselski et al. 2023). In liverworts, electrical signals have been reported to propagate at a rate of 0.4-1.5 mm/s (Trębacz et al. 1994, Koselski et al. 2021), while the dynamics of intracellular Ca^2+^ has not yet been explored. Nonetheless, the molecular mechanism underlying long-distance signal propagation in the absence of vascular bundles remain enigmatic.

*Marchantia polymorpha* holds particular significance due to several characteristics that render it suitable for investigating these molecular mechanisms. Notably, it is relatively amenable to genetic transformation, with the haploid generation encompassing most of its life cycle. Furthermore, the complete sequencing of the *Marchantia* genome has been achieved, revealing a genome that, despite retaining a multitude of genes encoding essential signaling factors conserved among land plants, demonstrates reduced genetic redundancy when compared to other land plants (Bowman et al. 2017).

In the present study, we provide groundbreaking evidence for the presence of wound-induced rapid Ca^2+^ wave and electrical signal propagation mechanisms in the non-vascular plant, *Marchantia polymorpha*. We induced localized damage to the thallus as a stressor and observed that the propagation velocity of both Ca^2+^ waves and electrical signals in *Marchantia* was approximately 1-2 mm/s, a rate akin to what has been reported for vascular plants. Furthermore, Ca^2+^ wave and electrical signal propagation were tightly coupled. Genetic studies notably highlight the crucial role of the sole GLR but do not implicate either type 1 or type 2 TPCs in the long-distance propagation of the Ca^2+^ wave and electrical signals. This compelling evidence suggests that the long-distance signaling mechanism mediated GLRs is evolutionarily conserved in both vascular and non-vascular land plants.

## Results

### Propagation of wound-induced Ca^2+^ waves and electrical signals in *Marchantia* thallus

We employed plastochron 2 stage plants, in which the thallus had divided into four branches (Fig. S1, Solly et al. 2017). In the asexual reproduction cycle, a *Marchantia* gemma develops dichotomous branching thallus from the two distinct meristematic zones (Fig. S1, Solly et al. 2017). In this study, we designate the wounded branch as “branch 1”, while both branch 1 and branch 2 originate from the same meristematic zone of the gemma. In addition, “branch 3” and “Branch 4” refer to branches that arise from different meristematic zones than “branch 1” and "branch 2”. The central region of the thallus is referred to as the intermediate site.

To visualize Ca^2+^ dynamics in *Marchantia*, we introduced a fusion protein consisting of GCaMP6f, a Ca^2+^ sensor, and mCherry, a Ca^2+^-independent fluorescent protein (Waadt et al. 2017). The fluorescence intensity of GCaMP6f and mCherry expressed with the Mp*EF1α* promoter was particularly high in the notch area and gemma cup (Fig. S2). This is consistent with the expression pattern of Mp*EF1α*. (Sauret-Güeto et al. 2020). Variations in fluorescence intensity from site to site are likely due to variations in the expression levels of the fluorescent protein. When a branch of thallus was locally wounded by a needle, increases in GCaMP6f fluorescence intensity were induced in both the wounded and unwounded sites (Fig. S2, Video S1). In contrast, mCherry fluorescence intensity did not change before or after wounding (Fig. S2). To better visualize increased fluorescence intensity in areas of low GCaMP6f expression, binarized images were created showing areas of significantly increased GCaMP6f fluorescence intensity in black, compared to pre-wounding levels. The [Ca^2+^]_cyt_ elevation propagated distally from the wounded site in a wave-like manner (Fig. 1A, B).

**Figure 1.**
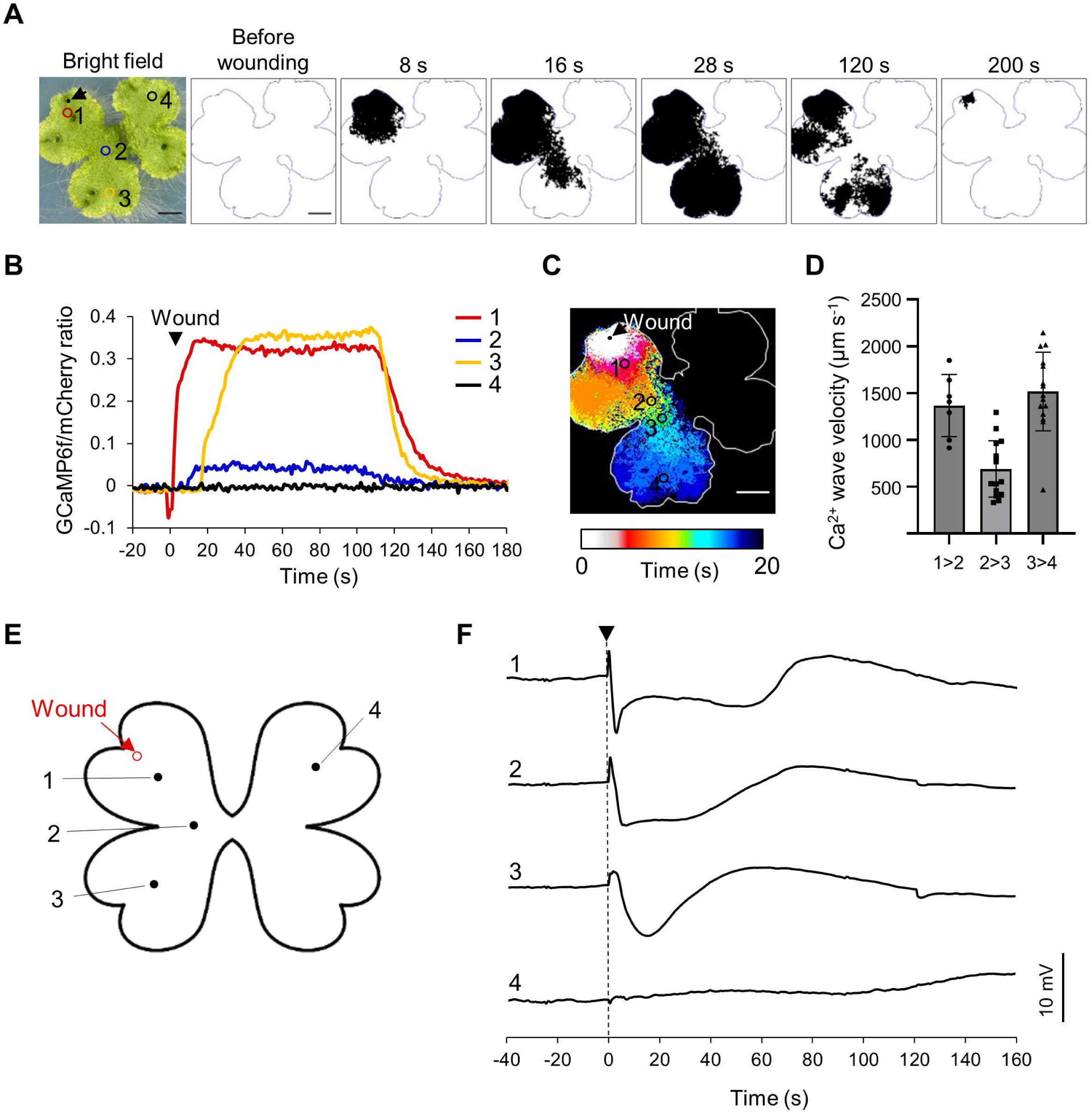
Wound-induced Ca^2+^ wave and electrical signal propagation in *Marchantia polymorpha*. (A) Bright field image and time lapse imaging of cytosolic Ca^2+^ in *Marchantia* thali expressing fluorescent proteins GCaMP6f-mCherry after wounding with a needle. Binarized images were created with areas of significantly increased fluorescence intensity filled in black. The standard for significantly increasing the fluorescence intensity was F > F_0_ + 2*SD (standard deviation). F_0_ = average fluorescence intensity of 10 frames before wounding. The arrow indicates wounded site. Scale Bar = 3 mm. (B) Measurement of GCaMP6f/mCherry ratio at positions 1–5 (ROI in image of A). Normalized by subtracting the average ratio value of the 10 frames before wounding from the ratio value at each time. The arrow indicates the time of wounding. (C) The time at which the Ca^2+^ wave arrived is shown. (D) Velocity of the Ca^2+^ wave between the positions 1 to 2, 2 to 3, and 3 to 4 in Figure C. n=7–12. mean ± SD. (E) Schematic diagram of the experiment. Surface potentials were measured at the wounded branch 1 (positions 1), at the intermediate site between branch 1 and 2 (position 2), at the unwounded branch 2 (position 3), and branch 3 (position 4). (F) Representative recording of wound-induced surface potential changes at positions 1–4. The triangular arrow indicates the time of wounding.

Wounding the branch 1 induced Ca^2+^ waves propagated within the two branches (branch and 2) emerging from the same meristematic zone of the gemmae in 92.8% (39/42) of the samples examined (Fig. S3A). The rest of the samples (7.2%; 3/42), the Ca^2+^ waves were confined only to the branch 1, with no propagation extending to the branch 2 (Fig. S3A). Interestingly, Ca^2+^ waves did not propagate to branch 3 and 4 beyond the intermediate site in all samples tested (Fig. 1, Fig. S1, Fig. S3A). The velocity of the Ca^2+^ waves, measured from positions 1 to 2, was 1512 ± 585 µm s^-1^ (n=8). However, this velocity decreased as the wave traversed between branches and subsequently accelerated upon reaching the interior of the branch (Fig. 1C, D).

To investigate whether electrical signal propagation is induced by wounding in *Marchantia*, we measured the surface potentials of the thallus at four positions. Electrode 1 was positioned in proximity to the wounded site, electrode 2 was positioned in the intermediate site between Branches 1 and 2, and electrode 3 was symmetrically placed with electrode 1 on Branch 2. Electrode 4 was situated on Branch 3. (Fig. 1E). Wounding induced changes in surface potentials in both wounded and unwounded branches (Fig. 1F). As with the Ca^2+^ waves, the change in surface potential did not propagate beyond the intermediate site to the two opposite branches (Fig. 1F). An identical response was observed in all 8 samples tested.

The initial positive change in surface potential is not dependent on the alteration of membrane potential due to the influx and efflux of ions directly beneath the measuring electrode. The measuring electrodes are electrically interconnected through the electrolyte within the apoplast and medium, establishing an electrical circuit. Consequently, a shift in surface potential at the stimulation site is detected as a positive surface potential change at an electrode distant from the stimulation site through the electrical circuit (Zawadzki and Trębacz 1985). Therefore, we focused on the timing of the decrease in surface potential. The decrease in surface potential was induced sequentially from position 1 to 3, providing clear evidence that mechanical injury triggers the propagation of electrical signals in *Marchantia*. Even with an increased number of electrodes aimed at examining the dynamics of the electrical signals in more detail, a decrease in surface potential was induced in the order of proximity to the wounded site (Fig. S4). Notably, the velocity of this electrical signal measured from positions 1 to 2, was 1517 ± 958 µm s^-1^ (n=8), a rate nearly identical to the propagation velocity of Ca^2+^ waves in *Marchantia* and electrical signals in vascular plants.

### Wound-induced [Ca^2+^]_cyt_ elevation and surface potential changes are linked

To elucidate the connection between electrical signals and Ca^2+^ waves, we conducted concurrent measurements of surface potential and [Ca^2+^]_cyt_ at two distal positions, surface potential at Electrode 1 (E1) – Electrode 2 (E2) and [Ca^2+^]_cyt_ at region of interest 1 (ROI1) - ROI2 (Fig. 2A). We set ROIs in close proximity to the electrodes since the fluorescence of GCaMP6f cannot be observed directly beneath the electrodes. To assess whether there is a discrepancy in the timing of the increase in GCaMP6f fluorescence intensity based on the ROI’s location near the electrode, we measured the fluorescence intensity of GCaMP6f in ROIs at various positions near the electrode (Fig. S6). There was no significant difference in the timing of the increase in GCaMP6f fluorescence intensity when ROIs were set at different positions in the vicinity of the electrode (Fig. S6). We presumed that, under the current measurement conditions, the placement of the ROI close to the electrode would not significantly affect the measurement of signal arrival timing. Therefore, we arbitrarily positioned the ROI near the electrode and conducted the measurements. The negative changes in surface potential resulting from mechanical injury and the elevation in [Ca^2+^]_cyt_ occurred almost simultaneously (Fig. 2A). To ascertain whether the electrical signal and the Ca^2+^ wave exhibit concurrent arrival, we assessed the latency time (time 1) from the moment of wounding to the occurrence of negative surface potential changes and the elevation in [Ca^2+^]_cyt_ at positions 1 and 2. Our analysis indicates that the latency between the [Ca^2+^]_cyt_ elevation and the negative surface potential changes is closely matched in value. (Fig. 2B). A correlation was identified in the duration time (time 2) of these two signals (Fig. 2C). Furthermore, in order to confirm the concurrent propagation of the electrical signal and the Ca^2+^ wave, we assessed the propagation velocity of the two signals from position 1 to position 2 in each plant (Fig. 2D). To determine the equivalence of the velocity of the Ca^2+^ wave and the electrical signal, we calculated the differences between their propagation velocity in each plant. The analysis demonstrated that the difference in velocity between the two signals was statistically indistinguishable from zero (Fig. 2E). Collectively, these findings strongly suggest an association between negative surface potential changes and [Ca^2+^]_cyt_ elevation.

**Figure 2.**
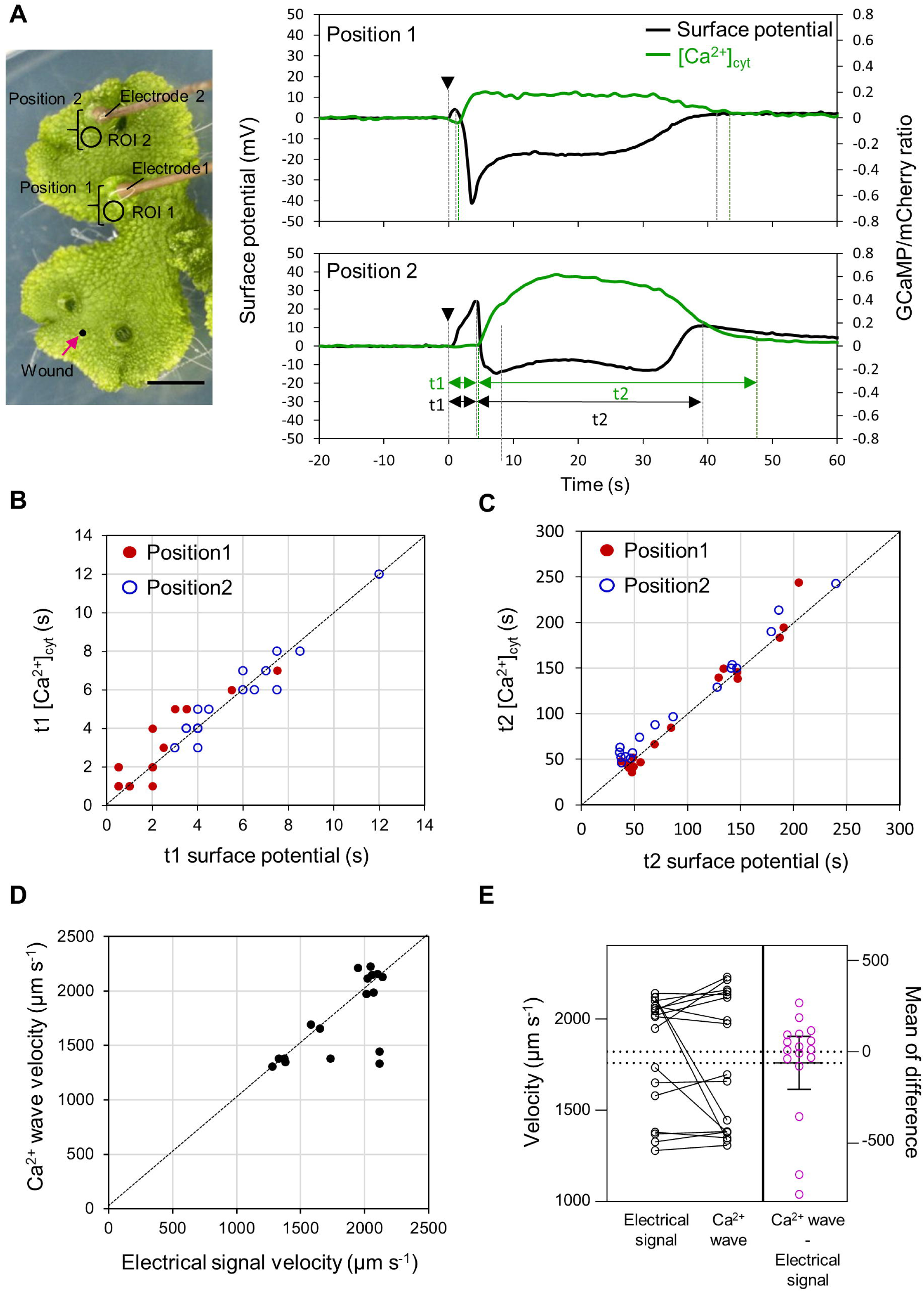
Simultaneous measurement of cytosolic Ca^2+^ concentration and surface potential. (A) Representative recording of fluorescence intensity of GCaMP6f at ROI1, ROI2 and surface potential at electrode 1, electrode 2. The triangular arrow indicates the time of wounding. Scale Bar = 3 mm. (B) Measurements of time 1 (time taken to induce a decrease in surface potential and an increase in GCaMP6f fluorescence intensity from the time of wounding) at position 1 and 2. (C) Measurements of time 2 (duration time of negative surface potential change and [Ca^2+^ ]_cyt_ elevation) at position 1 and 2. (D) Velocity of propagation of electrical signals and Ca^2+^ wave between position 1 and 2. (E) Comparison of velocity of the electrical signal and Ca^2+^ wave propagation. On the left side of the diagram, the rates of Ca^2+^ wave and electrical signal in each plant are plotted and connected by straight lines. The difference in the velocity of the two signals for each plant is plotted in the right side of the diagram. Mean ± 95% confidence interval. The 95% confidence interval includes 0, indicating that the difference in velocity between the two signals is statistically equal to 0. n = 17.

After wounding, positive surface potential change was induced preceding the [Ca^2+^]_cyt_ elevation at position 4 (Fig. 2A). This positive surface potential change was also induced at the site where the negative surface potential change and [Ca^2+^]_cyt_ elevation did not occur (Fig. S3). Positive change in surface potential after wounding appears to be independent of the [Ca^2+^]_cyt_ elevation and the negative surface potential changes.

### Critical importance of Ca^2+^ channels and K^+^ channels in the propagation of electrical signals and Ca^2+^ waves

To understand which ion channels are involved in Ca^2+^ wave and electrical signal propagation, we treated *Marchantia* with LaCl_3_, a Ca^2+^ channel inhibitor, and tetraethylammonium chloride (TEA), a potassium channel inhibitor.

When measuring surface potential, the plants were placed in plastic chambers divided into multiple sections, and the section containing the inhibitor was partially treated by sealing it with Vaseline as a barrier to prevent its diffusion into the other sections. Surface potentials were measured in the inhibitor-treated section.

In Ca^2+^ imaging, the observation of plants within a plastic enclosure posed challenges. Therefore, inhibitor treatment was administered by immersing the plant in a solution containing the inhibitor. Immersion in the solution led to an elevated percentage of individuals in which the Ca^2+^ waves were restricted to propagation solely within the wounded branches. This phenomenon was observed in 36.3% of the tested individuals, even when they were immersed in the solution without the inhibitor.

LaCl_3_ inhibited both Ca^2+^ wave and electrical signal propagation (Fig. 3A-C, Video S2), suggesting that Ca^2+^ influx from the apoplast through the plasma membrane is responsible for [Ca^2+^]_cyt_ elevation and membrane potential depolarization. Propagation of Ca^2+^ waves necessitate the propagation of electrical signals.

**Figure 3.**
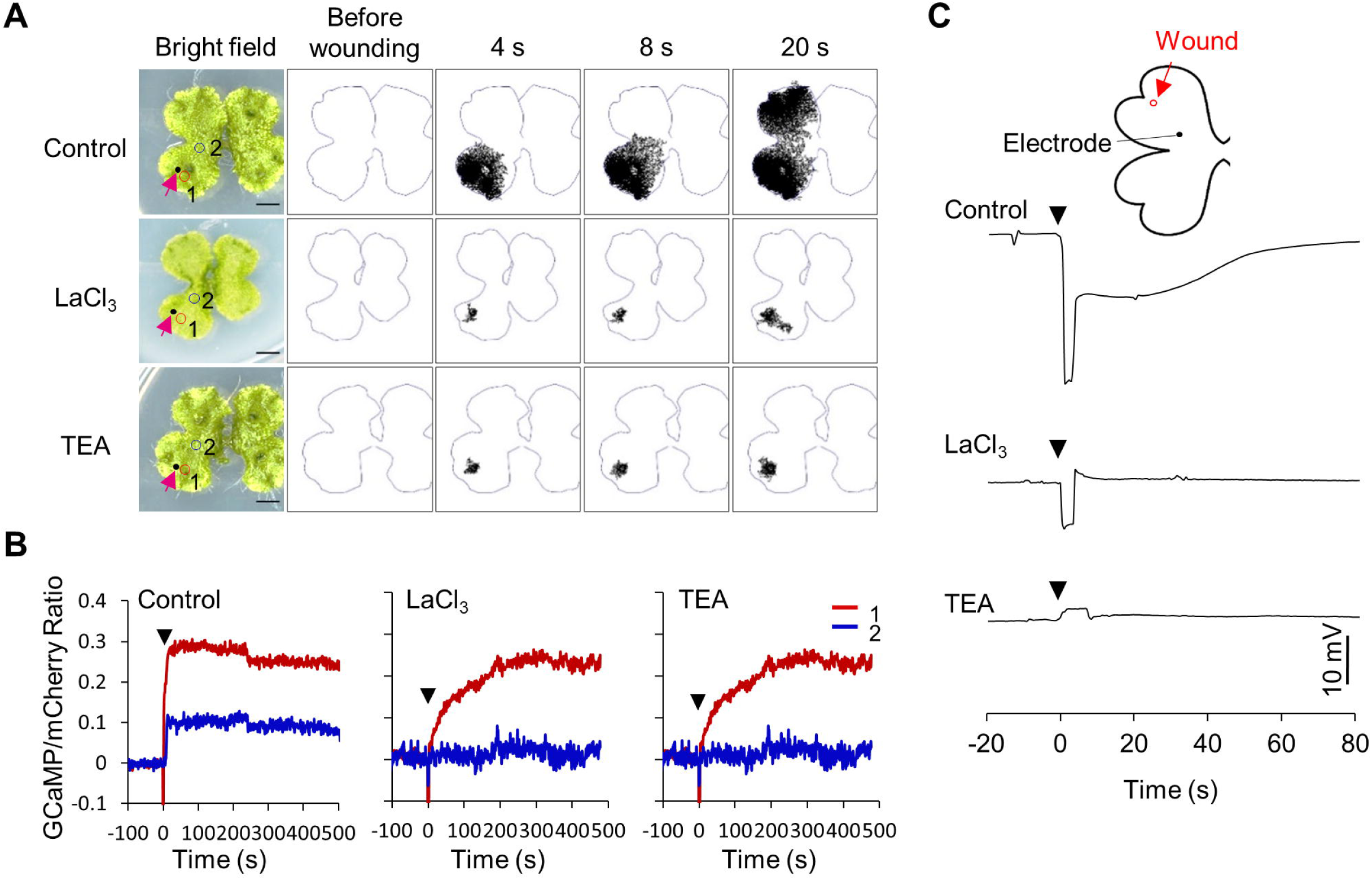
Effects of Ca^2+^ channel and K^+^ channel inhibitors on wound-induced Ca^2+^ wave and electrical signal propagation. (A) Binarized time lapse imaging of cytosolic Ca^2+^ in *Marchantia* expressing GCaMP6f-mCherry after wounding. Plants were pretreated with LaCl_3_ (5 mM) or tetraethylammonium chloride (TEA; 5 mM) for 30 min. The magenta arrows indicate wounded site. Scale Bar = 3 mm. (B) Representative recording of GCaMP6f/mCherry ratio value in position 1, 2. (C) Measurement of surface potential after wounding. The triangular arrows indicate the time of wounding.

Despite the critical importance of Ca^2+^ channels, possible involvement of K^+^ channels have remained poorly understood. TEA also suppressed both electrical signals and Ca^2+^ waves, suggesting the importance of K^+^ channels in the long-distance propagation of Ca^2+^ waves and electrical signals.

### MpGLR is required for rapid long-distance signal propagation

*Marchantia polymorpha* has a single GLR that is phylogenetically close to the clade3 GLRs in *Arabidopsis thaliana* (Wudick et al. 2017). We have generated Mp*glr-1^ko^* by genome editing and investigated whether MpGLR is required to propagate wound-induced Ca^2+^ waves and electrical signals (Fig. S7, AB). In Mp*glr-1^ko^*, wound-induced Ca^2+^ wave and electrical signal propagation were entirely suppressed (Fig. 4A, B, Video S3). On the other hand, the [Ca^2+^]_cyt_ elevation and surface potential changes induced at the local wounded site was not suppressed in Mp*glr-1^ko^* (Fig.4C, D). In addition, slow Ca^2+^ wave propagation was observed at wounded sites in Mp*glr-1^ko^* (Fig. 4C, Video S4). To further validate whether GLR is essential for signal propagation, we developed a line in which *proEF1α:*Mp*GLR* was introduced into the Mp*glr-1^ko^* mutant. The incorporation of *GLR* complementary DNA into the plant body was confirmed by conducting PCR with primers designed to amplify the intron-containing region (Fig. S7A, C). Subsequently, after generating the complementation line, GCaMP6f was introduced, resulting in different expression levels of GCaMP6f in the Tak-1 and GLR complemented lines. Wound-induced Ca^2+^ wave and electrical signal propagation were observed in the complementation line (Fig. 4A, Video S3). Similar to the wild type (WT), Ca^2+^ waves did not propagate beyond the intermediate site to Branches 3 and 4 in the complementation line. The velocity of signal propagation seemed to be faster in the complementation line than in the WT, possibly due to the expression of Mp*GLR* by the *EF1α* promoter. In all of the eight tested complementation lines, Ca^2+^ waves propagated throughout Branch 1 and Branch 2 within one or two frames immediately after wounding. Consequently, accurate measurement the velocity of signal propagation in the complementation line was not feasible. These results strongly suggest that MpGLR plays a crucial role in modulating the propagation of wound-induced Ca^2+^ waves and electrical signals. However, other ion channels, distinct from MpGLR, are likely responsible for [Ca^2+^]_cyt_ elevation and potential surface changes at the local wounded site.

**Figure 4.**
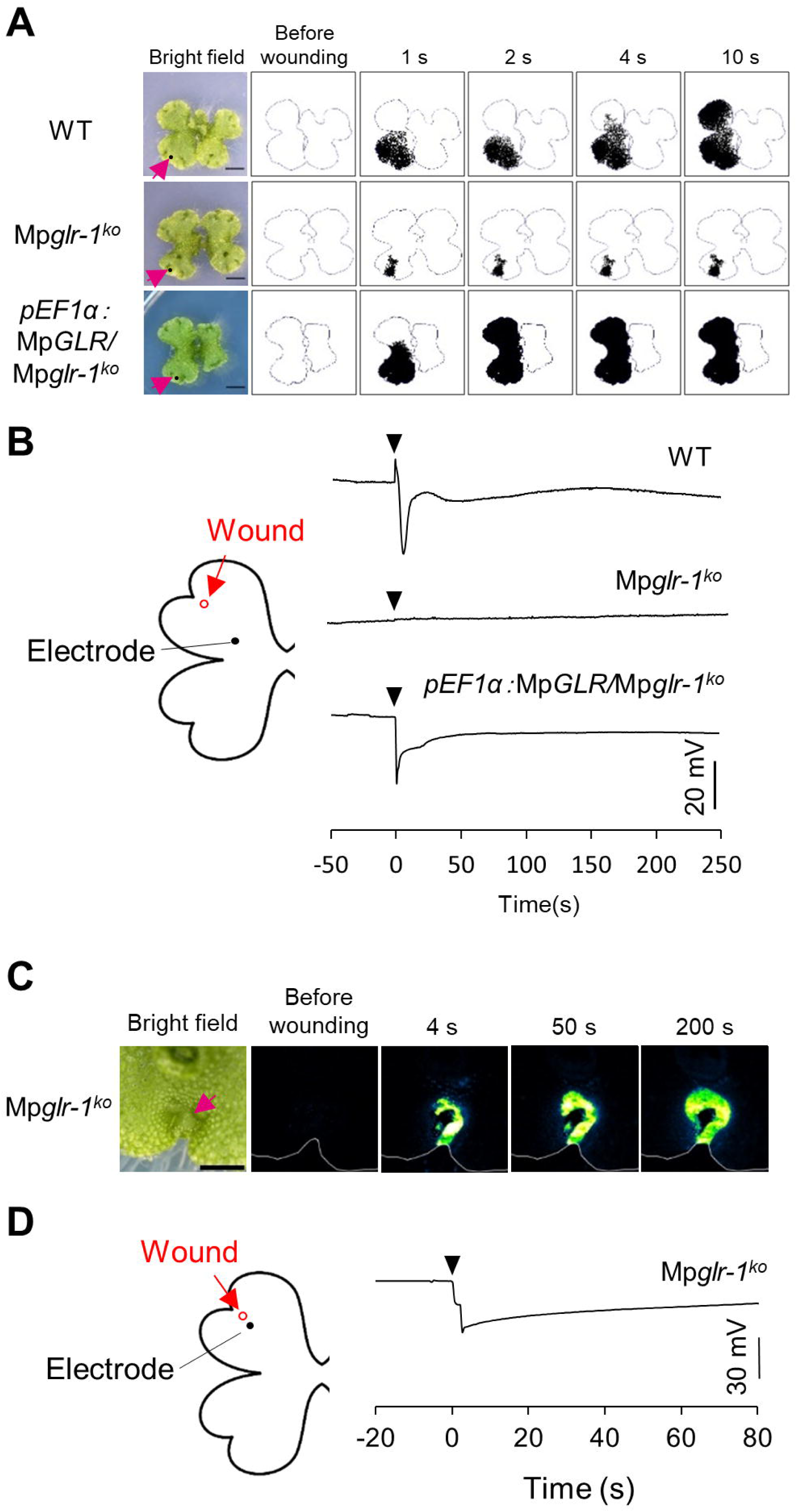
Mp*GLR* is essential for electrical signal and Ca^2+^ wave propagation. (A) Time lapse imaging of cytosolic Ca^2+^ in WT, Mp*glr-1^ko^*, *pEF1α:*Mp*GLR/*Mp*glr-1^ko^*(complementation) line expressing GCaMP6f-mCherry after wounding. The magenta arrows indicate wounded site. Scale Bar = 3 mm. (B) Representative recording of surface potential in the wild type (WT), Mp*glr-1^ko^* and *pEF1α:*Mp*GLR/*Mp*glr-1^ko^*(complementation) line after wounding. (C) Time lapse imaging of cytosolic Ca^2+^ at the wounded site in Mp*glr-1^ko^*. Scale Bar = 1.5 mm. (D) Representative recording of surface potential at wounded site in Mp*glr-1^ko^*.

### MpTPCs are not required for long-distance signal propagation

Since TPC has been reported to be required for Ca^2+^ wave propagation in *Arabidopsis* (Choi et al. 2014, Kiep et al. 2015), we investigated whether MpTPCs are involved in Ca^2+^ wave and electrical signal propagation. *Marchantia polymorpha* possesses three homologs of Two-Pore Channel (TPC) proteins, which are localized at the vacuolar membrane. Among these, only MpTPC1 exhibits Slow Vacuolar (SV) channel activity, akin to the TPCs found in vascular plants. In contrast, MpTPC2 and MpTPC3 belong to a distinct group separate from the TPCs found in vascular plants and notably lack SV channel activity (Hashimoto et al. 2022).

We examined the electrical signals and [Ca^2+^]_cyt_ kinetics in the mutants defective in Mp*TPC1* and Mp*TPC2/MpTPC3* (Hashimoto et al. 2022). In Mp*tpc1-1^ko^* and Mp*tpc2-2 tpc3-2^ko^*double mutant, wound-induced Ca^2+^ wave propagation was observed as in the wild type (WT) (Fig. 5A, Video S5). No significant differences were observed in the signal propagation velocity (Fig. 5B). Wound-induced electrical signals also propagated at a comparable velocity to WT (Fig. 5C). These results suggest that the MpTPCs are not involved in Ca^2+^ wave and electrical signal propagation.

**Figure 5.**
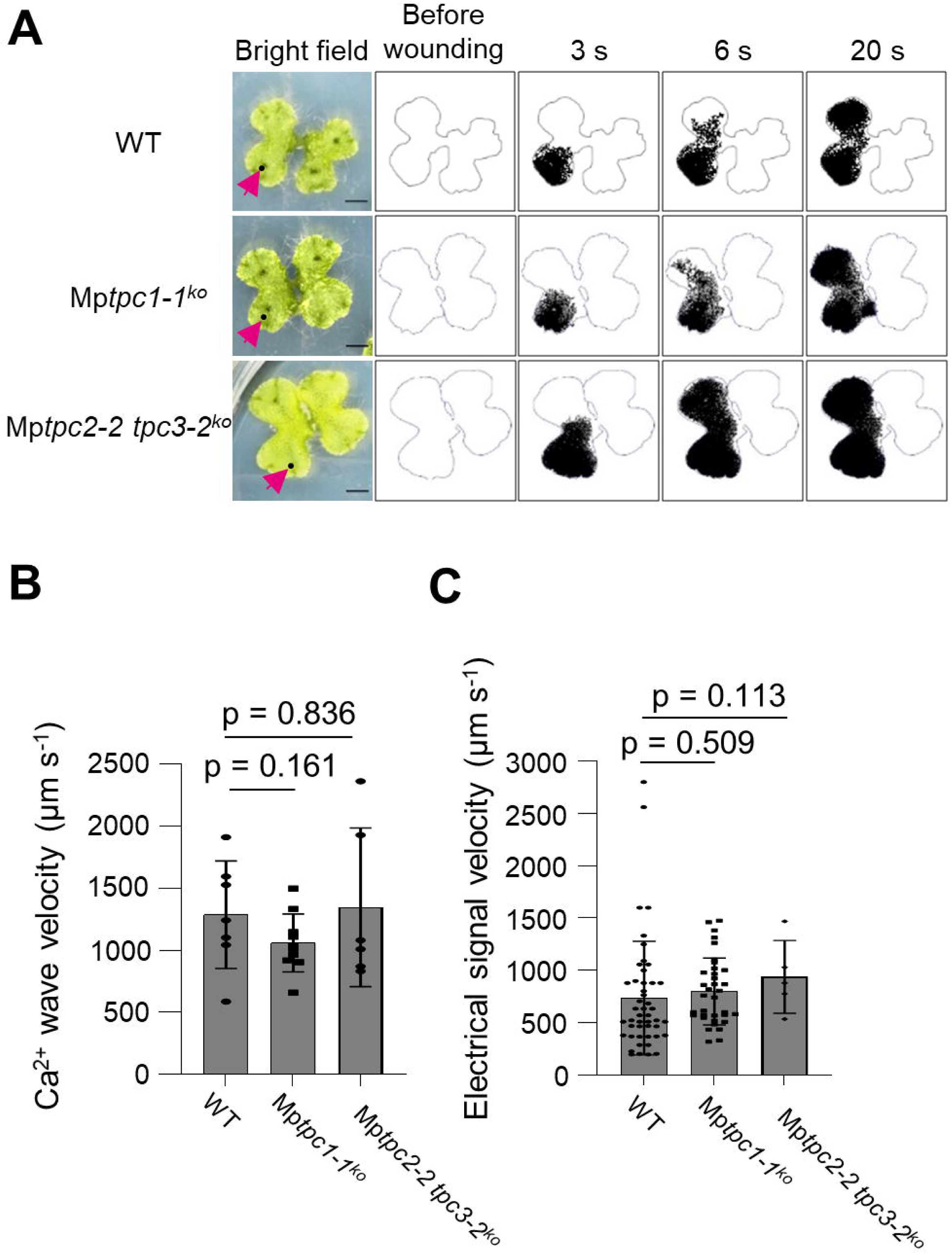
None of the three Mp*TPCs* are required for electrical signal and Ca^2+^ wave propagation. (A) Binarized time lapse imaging of cytosolic Ca^2+^ in WT and Mp*tpc1-1^ko^*, Mp*tpc2-2 tpc3-2^ko^* expressing GCaMP6f-mCherry after wounding. The magenta arrow indicates the wounded site. Scale bar = 3 mm. (B) Velocity of Ca^2+^ wave in WT, Mp*tpc1-1^ko^* and Mp*tpc2-tpc3-2^ko^*. n=7-10. mean ± SD. (C) Velocity of electrical signal in WT, Mp*tpc1-1^ko^* and Mp*tpc2-2 tpc3-2^ko^*. n = 5-35. P-values were calculated by Mann-Whitney test.

## Discussion

Vascular plants propagate long-distance Ca^2+^ and electrical signals from localized stress sites to distant tissues through the vascular bundle. Various models have recently been proposed for the mechanisms underlying long-distance signaling, grounded in the presence of vascular bundles (Farmer et al. 2014, Bellandi et al. 2022, Gao et al. 2023, Grenzi et al. 2023). However, in the present study, we demonstrated that *Marchantia polymorpha* lacking vasculature exhibits a wound-induced rapid long-distance signaling. In this signaling paradigm, Ca^2+^ waves and electrical signal propagation are intricately intertwined. Remarkably, we observed that the propagation velocity of both Ca^2+^ waves and electrical signals across the thallus was approximately 1-2 mm/s (Fig. 1-2), a rate reminiscent of what is observed in the vascular bundles of angiosperms (Mousavi et al. 2013, Nguyen et al. 2018, Toyota et al. 2018). The propagation of these two signals was also suppressed in Mp*glr-1^ko^* mutant (Fig. 4), suggesting that a rapid long-distance signaling mechanism involving GLR is conserved in *Marchantia*.

In recent years, several models have been considered for the mechanism of long-distance signal propagation. For example, a calcium-induced calcium release model in which [Ca^2+^]_cyt_ elevation itself induces [Ca^2+^]_cyt_ elevation in adjacent cells has been proposed (Evans et al. 2016, Choi et al. 2017). On the other hand, some models do not require [Ca^2+^]_cyt_ elevation or depolarization of the plasma membrane potential for signal propagation. One model is that wound-induced change in turgor pressure in the vascular bundle propagates (hydraulic signal), and the pressure change activates the GLR by unknown mechanism, inducing [Ca^2+^]_cyt_ and depolarization of the plasma membrane potential (Farmer et al. 2014, Grenzi et al. 2023). Another model is that substances that activate GLR and induce [Ca^2+^]_cyt_ elevation and depolarization of the membrane potential diffuse thorough vascular bundle from the wounded sites (Bellandi et al. 2022, Gao et al. 2023). In this study, we have shown that a rapid Ca^2+^ wave and electrical signal propagation mechanism involving GLR is conserved in non-vascular plants (Fig.4). When the signal propagated from the wounded Branch 1 to Branch 2, its propagation velocity decelerated between the two Branches and subsequently accelerated (Fig.1C, D). These results imply that rapid Ca^2+^ wave and electrical signal propagation in *Marchantia* cannot be attributed to the direct diffusion of substances that inducing plasma membrane depolarization or [Ca^2+^]_cyt_ elevation from the wounded sites to distal sites. Whether GLRs can be activated by alterations in plasma membrane potential or by an increased concentration of glutamate in the apoplast has been a topic of ongoing debate. (Moe-Lange et al. 2021, Matteo et al. 2023). Testing these possibilities may lead to understanding of the mechanism of GLR activation in non-vascular plants.

The potential interconnection and reciprocal regulation between Ca^2+^ waves and electrical signals have been a topic of discussion (Gilroy et al. 2016, Choi et al. 2017). Ca^2+^ has been suggested to be involved in membrane depolarization in various plants (Vodeneev et al. 2016). Inhibition of Ca^2+^ influx from outside the cell by La^3+^ suppressed wound-induced electrical signal propagation in *Marchantia* (Fig. 3). Both wound-induced Ca^2+^ waves and electrical signal propagation depend on Mp*GLR* (Figs. 4). These results indicate that GLR-mediated rapid long-distance signaling involving Ca^2+^ waves and electrical signals appears to be highly conserved among vascular and non-vascular land plants.

[Ca^2+^]_cyt_ elevation and surface potential changes induced at the local wounded site was not suppressed in Mp*glr-1^ko^* or La^3+^, TEA treatment (Fig. 3, 4), and slow Ca^2+^ wave propagation was observed at wounded sites in Mp*glr-1^ko^* (Fig. 4B, Video S4). In the vicinity of the wounded sites, other ion channels, distinct from MpGLR, may play a crucial role in local plasma membrane depolarization and propagation of slow and localized Ca^2+^ wave. The slow and localized Ca^2+^ wave may also be attributed to the diffusion of Ca^2+^ from the wounded cells into the surrounding cells. Given the much less genetic redundancy in *Marchantia polymorpha* in comparison with angiosperms such as *Arabidopsis thaliana* (for example, only 1 GLR in *Marchantia*, while20 GLRs in *Arabidopsis*), the present experimental system would provide an excellent model to identify various ion channels involved in and elucidate the molecular mechanisms for rapid long-distance signaling conserved in land plants.

The long-distance signal propagation was inhibited not only by a Ca^2+^ channel blocker but also by TEA, an inhibitor for K^+^ channels (Fig. 3). TEA inhibits depolarization of cell membrane potential in *Conocephalum conicum* and *Physcomitrium* (*Physcomitrella) patens* (Trebacz et al. 1989, Johannes et al. 1997). In *Conocephalum*, the equilibrium potential of K^+^ is higher than the resting membrane potential, opening of the K^+^ channel may induce depolarization of the membrane potential (Trebacz et al. 1994, Kisnieriene et al. 2022). Furthermore, simultaneous measurement of [Ca^2+^]_cyt_ and surface potential revealed that these two signals are tightly interconnected (Fig. 2). These results underscore the critical role of K^+^ channels in mediating membrane depolarization, and both [Ca^2+^]_cyt_ elevation and depolarization of the plasma membrane potential are required for the propagation of these two signals.

Wound-induced Ca^2+^ waves and electrical signals failed to propagate to half of the body through intermediate sites (Fig. 1). *Marchantia* exhibits a reproductive strategy involving the formation of clones known as gemmae, which proliferate within specialized structures called gemmae cups during asexual reproduction. Gemmae remain in a dormant state characterized by arrested cell division and differentiation. Notably, gemmae possess two apical meristems, which become active upon breaking dormancy, initiating growth from the apical meristems (Shimamura, 2015). The cells located in the intermediate sites where Ca^2+^ waves fail to propagate are presumed to be gemmae cells by origin. These gemmae cells differ from other thallus cells in that they have remained in a dormant state and did not originate from the apical meristem. The present study has demonstrated that the sole MpGLR is crucial for the propagation of Ca^2+^ waves and electrical signals (Fig. 4). Additionally, studies involving the overexpression of PLASMODESMATA-LOCATED PROTEIN 5 (PDLP5) and the examination of plasmodesmata (PD)-associated β-1,3-glucanase-deficient mutants with reduced PD conductance have indicated that Ca^2+^ wave propagation at non-vascular sites is inhibited (Toyota et al., 2018). It is plausible that gemmae cells have lower expression levels of factors associated with rapid long-distance signaling. Consequently, this reduced expression may account for the failure of Ca^2+^ waves to propagate in gemmae cells.

Ca^2+^ wave and electrical signal propagation was induced in Mp*tpc1^ko^* mutants as in WT (Fig. 5). In *Arabidopsis tpc1* mutant, wound-induced systemic [Ca^2+^]_cyt_ elevation is suppressed at distal sites (Kiep et al. 2015). TPC opening is facilitated by Ca^2+^ binding to the EF-hand at the cytosolic (Beyhl et al. 2009, Schulze et al. 2011). It has been proposed that Two-Pore Channels (TPCs) may be activated initially by a rise in Ca^2+^ levels, potentially regulated by factors such as GLRs, and subsequently amplify the signal by modulating Ca^2+^ release from the vacuole (Kiep et al., 2015). Conversely, recent studies have reported the induction of Ca^2+^ wave and electrical signal propagation in *tpc1* mutants (Moe-Lange et al. 2021, Bellandi et al. 2022). In *Marchantia*, neither MpTPC1 (type 1 TPC) nor MpTPC2 and MpTPC3 (type 2 TPCs) seem to be indispensable for the propagation of the Ca^2+^ waves and electrical signals themselves.

In summary, this study demonstrates that GLR-mediated long-distance signaling mechanisms are conserved in liverworts, non-vascular plants, and presumably in all land plants. However, many questions remain unanswered regarding the activation of GLR during signal propagation and the factors that play a role in signal propagation itself. *Marchantia*, characterized by its relatively low genetic redundancy and the absence of vascular bundles, serves as a promising model organism for investigating the factors involved in long-distance signaling and gaining insights into the propagation mechanisms.

## Materials and Methods

### Plant material and growth conditions

*Marchantia polymorpha* accession Takaragaike-1 (Tak-1; male) were used as wild-type. Plants were grown on half-strength Gamborg’s B5 solid medium containing 1% agar, 1% sucrose, 1 g/L MES (pH 5.5, adjusted by KOH) under continuous light (approximately 40 µmol m^-2^s^-1^) at 21°C. 14-day-old plants were used for Ca^2+^ imaging and surface potential measurements.

### Plasmid construction and transformation

GCaMP6f-mCherry was cloned into the NotI-AscI site of pENTR/D-TOPO (Invitrogen). The entry clones were subcloned into pMpGWB303 vector (Ishizaki et al. 2015) using Gateway LR reaction. Complementary DNA (cDNA) of MpGLR was cloned into pENTR/D-TOPO using primers 5’-TTTTGCGGCCGCTCAAATGACACTATTTCTGTCC-3’ and 5’- TTTTGGCGCGCCTCAAATGACACTATTTCTGTCC-3’. The entry clones were subcloned into pMpGWB337 vector (Nishihama et al. 2016) using Gateway LR reaction. To obtain Mp*glr-1^ko^* mutant the gRNA were designed using a web tool CRISPRdirect (https://crispr.dbcls.jp) and cloned into the pMpGE013 vector (Sugano et al. 2018). These vectors were transformed into Agrobacterium GV2260. Transformation into *Marchantia* was performed using the G-Agar trap method, transformants were selected using hygromycin (Tsuboyama and Kodama 2018). To confirm that the MpGLR cDNA had been introduced, DNA was extracted from the plants and PCR was performed using two primers, 5’- AGCTCCACCAAGCCCAAGTC-3’ and 5’-CGCTTGGTGGTTATCGTTAC-3’.

### Wounding assay

A robotic arm, Do-bot MG400 (product model; DR-MG-4R005-02E), was equipped with a 20 cm long 5 mm square plastic rod. A needle (tungsten, 0.5 mm diameter) was attached to the end of the rod. Wounding was inflicted by the robotic arm, which was inserted vertically just above the thallus so that the needle penetrated the thallus completely.

### Ca^2+^ imaging

For Ca^2+^ imaging, GCaMP6f-mCherry-expressing lines were used. The images were acquired by a Nikon SMZ25 equipped with Nikon DS-Ri2 camera, P2-SHR Plan Apochromatic 1X objective lens and Nikon INTENSILIGHT C-HGFIE lamp. For GCaMP6f image acquisition, a 470/40 nm excitation filter, a 500 nm long-pass dichroic, and a 535/50 nm emission filter were used, and for mCherry images acquisition, a 545/15 nm excitation filter, 570 nm long-pass dichroic, and 620/30 nm emission filter were used. To increase the number of images that could be acquired per unit time, fluorescence images of GCaMP6f and mCherry were not taken simultaneously, but only fluorescence images of GCaMP6f were taken to calculate Ca^2+^ wave velocity. In this case, fluorescence images of mCherry were taken for 20 seconds before acquiring images of GCaMP6f.

### Image analyses

Image analysis was performed using imageJ Fiji (Schindelin et al. 2012). Fluorescence intensity values were obtained as the average intensity within a circular ROI. GCaMP6f intensity were divided by mCherry intensity, and the calculated GCaMP6f/mCherry ratio was normalized by subtracting the average ratio values of the 10 frames before wounding from the ratio value at each time. The velocity was calculated by dividing the lag time for a significant fluorescence increase of GCaMP6f within each ROI by the distance between the ROIs. The time at which the fluorescence intensity of GCaMP6f exceeded the average fluorescence intensity + 3*SD of the 10 frames before wounding was considered the time at which a significant increase in fluorescence occurred. ΔF images and video s were created by subtracting the average image of the 10 frames before wounding from each frame. Binarized images were created by subtracting the average +2*SD image of the 10 flames before wounding from each frame. To calculate duration time, the derivative of GCaMP6f fluorescence intensity and surface potential was used. The duration time of [Ca^2+^]_cyt_ elevation was calculated as the time from when the derivative of the fluorescence intensity of GCaMP6f exceeded the average values + 3*SD of the 10 values before wounding, to the time when it dropped below this threshold. The duration time of surface potential was calculated as the time from when the derivative of surface potential dropped below the average values - 3*SD of the 10 values before wounding, to the time when it exceeded threshold.

### Surface potential measurement

Individual thalli together with agar were placed in small Petri dishes (35 mm). The Petri dish containing the thallus was covered with a stretched transparent foil to avoid evaporation. Thin sharp Ag/AgCl electrodes were gently inserted into the thallus through the foil, one day before the experiment. The reference electrode was placed on the other part of the plant, behind the narrowing through which neither Ca^2+^ waves nor electrical signals were able to pass. Electrical signals (surface potential changes) were registered with an iWorx 304T (iWorx, USA) amplifier running under a Labscribe 2.0 software (iWorx, USA).

For the concurrent measurement of [Ca^2+^]_cyt_ and surface potential, a custom-designed device was employed (Fig.S5). A 5 µL volume of 0.8% agarose gel containing 10 mM KCl was placed on the surface of thallus, through which an electrode was brought into contact. The reference electrode was placed in the medium. The gain of the device was set to 60x and connected to an Arduino Uno A/D converter, and the voltage was measured every 0.5 s. Arduino IDE is used to send/receive data and write programs to/from Arduino Uno.

### Inhibitor treatment

5 mM LaCl_3_ and 5 mM TEA were dissolved in 10 mM HEPES buffer (pH 5.5, adjusted by KOH). In surface potential measurements, the plants were placed in a plastic case divided into three compartments, one of which was filled with solution. The compartment containing the inhibitor was sealed with Vaseline to prevent the inhibitor from leaking into other compartments. The solution was applied for 30 minutes and then removed before measurements were started. In Ca^2+^ imaging, plants were completely immersed in the solution for 30 minutes. The solution was removed by aspiration, and the Ca^2+^ imaging was performed 2 hours later. In electrical signal measurement, elliptic well c.a. 3 -5 mm made of Vaseline was formed on one of the branches of the thallus. The well was filled with a standard solution supplemented with an ion channel inhibitor. Electrodes were placed inside and outside the wells. Stimuli were applied 30 minutes after filling the wells with the solution.

## Data Availability Statement

The data underlying this article are available in the article and in its online supplementary material.

## Funding

This work was supported in part by Grant-in-Aid for Scientific Research on Innovative Areas JP22H04734 from MEXT, Japan, JSPS KAKENHI Grant Number 23K02505 and Bilateral Joint Research Project #JPJSBP120214601.

## Disclosures

Conflicts of interest: No conflicts of interest declared.

## Acknowledgments

We thank Dr. Rainer Waadt for providing GCaMP6f-mCherry plasmid.

## Notes

### Competing Interest Statement

The authors have declared no competing interest.

### Summary of Updates

Figure 1, 3, 4, 5 revised. Supplemental Figure revised. We added one author.

