## Supplemental Figure for "Rapid propagation of Ca^2+^ waves and electrical signals in a liverwort *Marchantia polymorpha*"

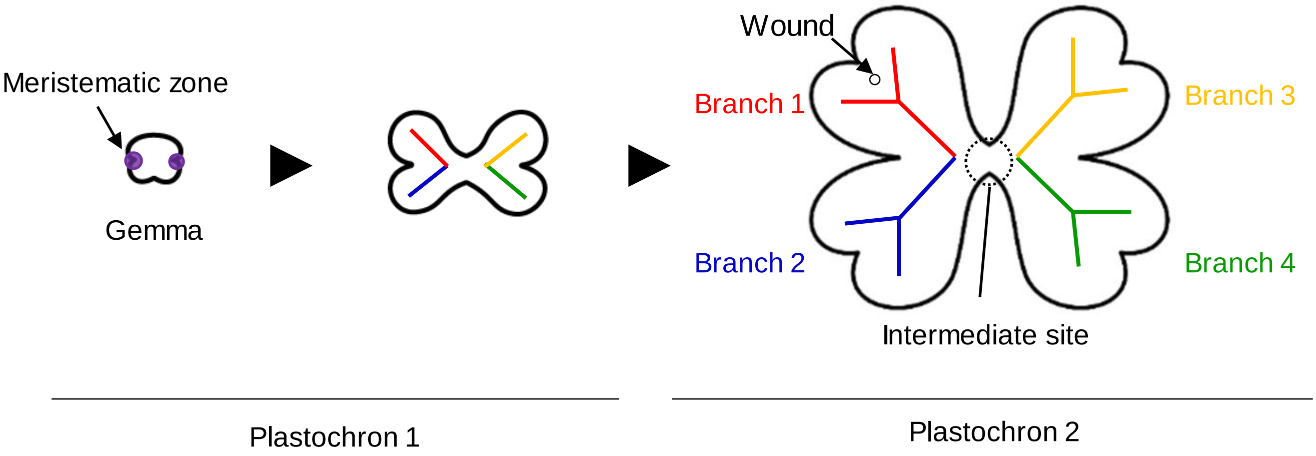


**Figure S1. Schematic diagram depicting the development of gemma and thallus in *Marchantia polymorpha*.** In the present research, we basically used plants at the growth stage with four branches. The wounded branch was designated as Branch 1. It is noteworthy that Branch 1 and Branch 2 (as well as Branch 3 and Branch 4) originate from the same meristematic zone of gemma.

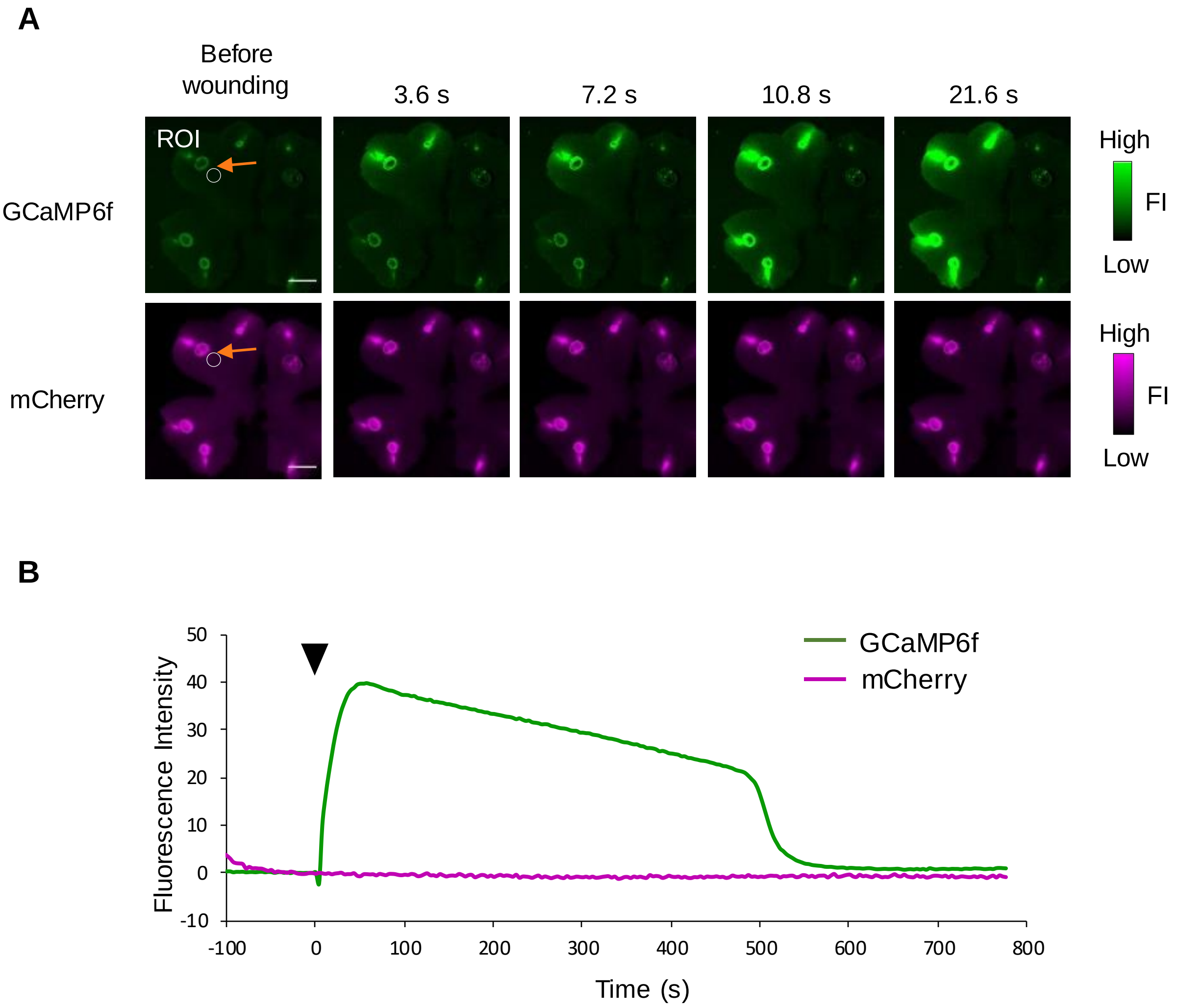


**Figure S2. Wound-induced $Ca^{2+}$ wave propagation in *Marchantia polymorpha*.** (A) Time lapse imaging of cytosolic $Ca^{2+}$ in *Marchantia* expressing GCaMP6f-mCherry after wounding. Orange arrows indicates the wounded site. Scale Bar = 3 mm. (B) Representative measurements of fluorescence intensity for GCaMP6f and mCherry in the region of interest (ROI). The triangular arrow indicates the time of wounding.

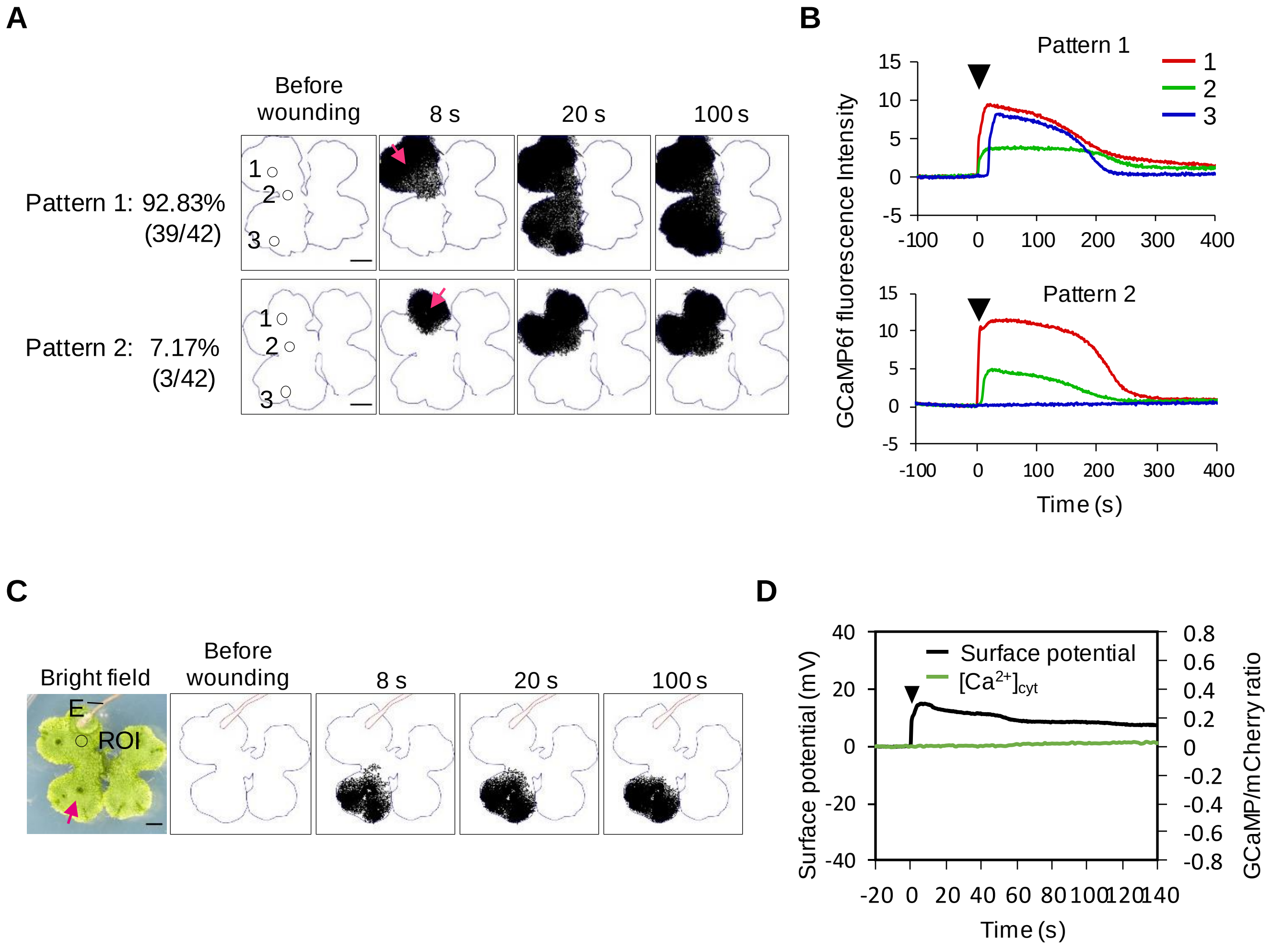


**Figure S3. Patterns of wound-induced $Ca^{2+}$ wave propagation.** (A and C) Binarized time lapse imaging of cytosolic $Ca^{2+}$ in *Marchantia* expressing GCaMP6f-mCherry after wounding. Magneta arrows indicates the wounded site. E represents the position of the electrode. Scale Bar = 3 mm. (B) Fluorescence intensity of GCaMP6f in position 1-3. (D) Simultaneous measurement of surface potential and GCaMP6f/mCherry ratio at electrode and ROI. The triangular arrow indicates the time of wounding.

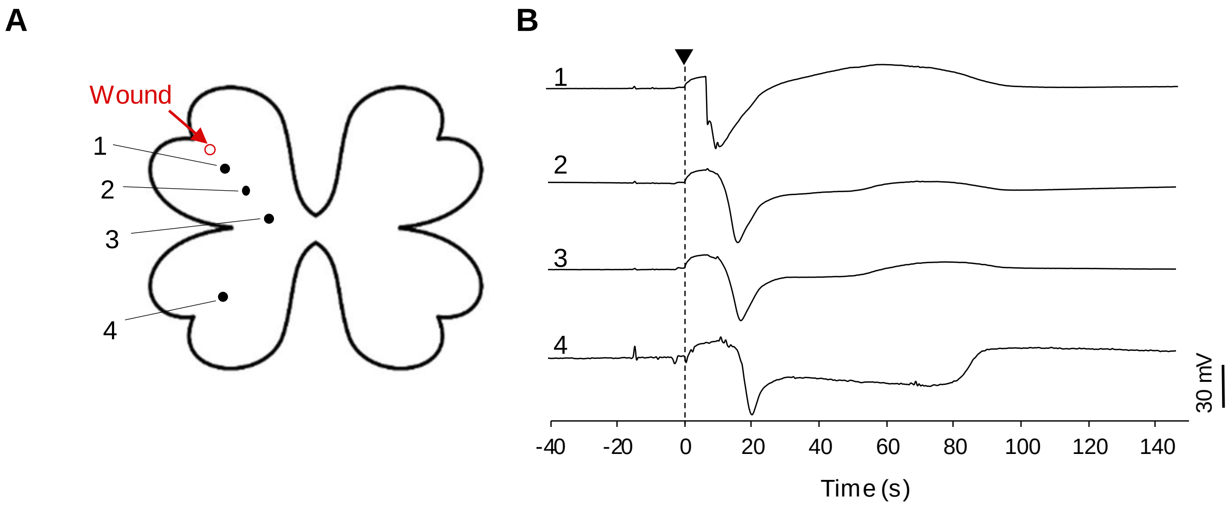


**Figure S4. Wound-induced electrical signal propagation.** (A) Schematic diagram of the experiment. Surface potentials were measured within the wounded branch (positions 1 and 2), at the intermediate site between the Branch 1 and Branch 2 (position 3), and within the Branch 2 (position 4). (B) Representative recording of wound-induced surface potential changes at positions 1 – 4. The triangular arrow indicates the time of wounding.

**A**

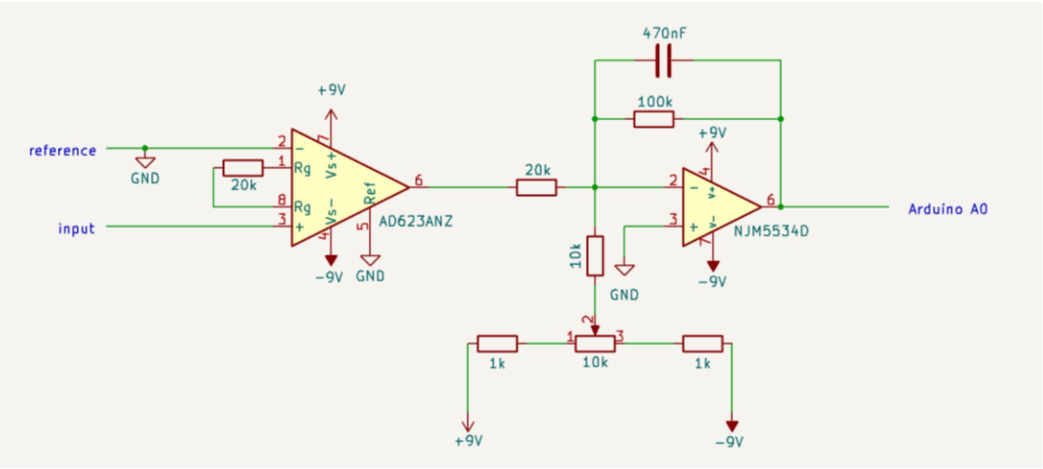


**B**

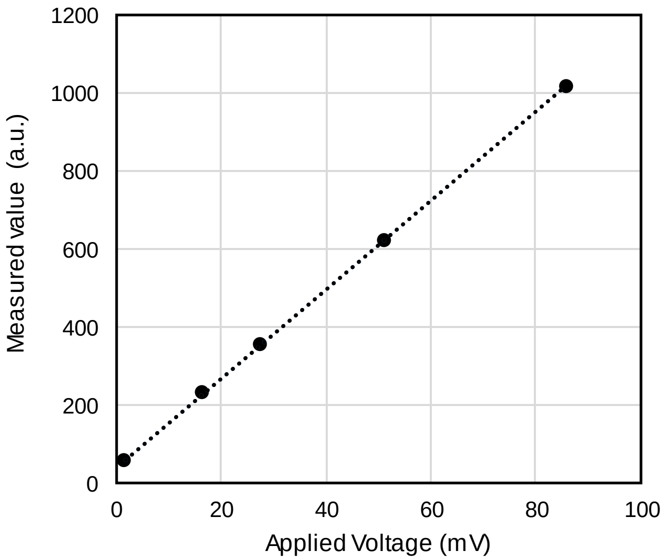


**Figure S5 The equipment to measure the surface potential.** (A) Circuit diagram depicting a self-made device designed for surface potential measurement. (B) Assessment of voltage measurement linearity.

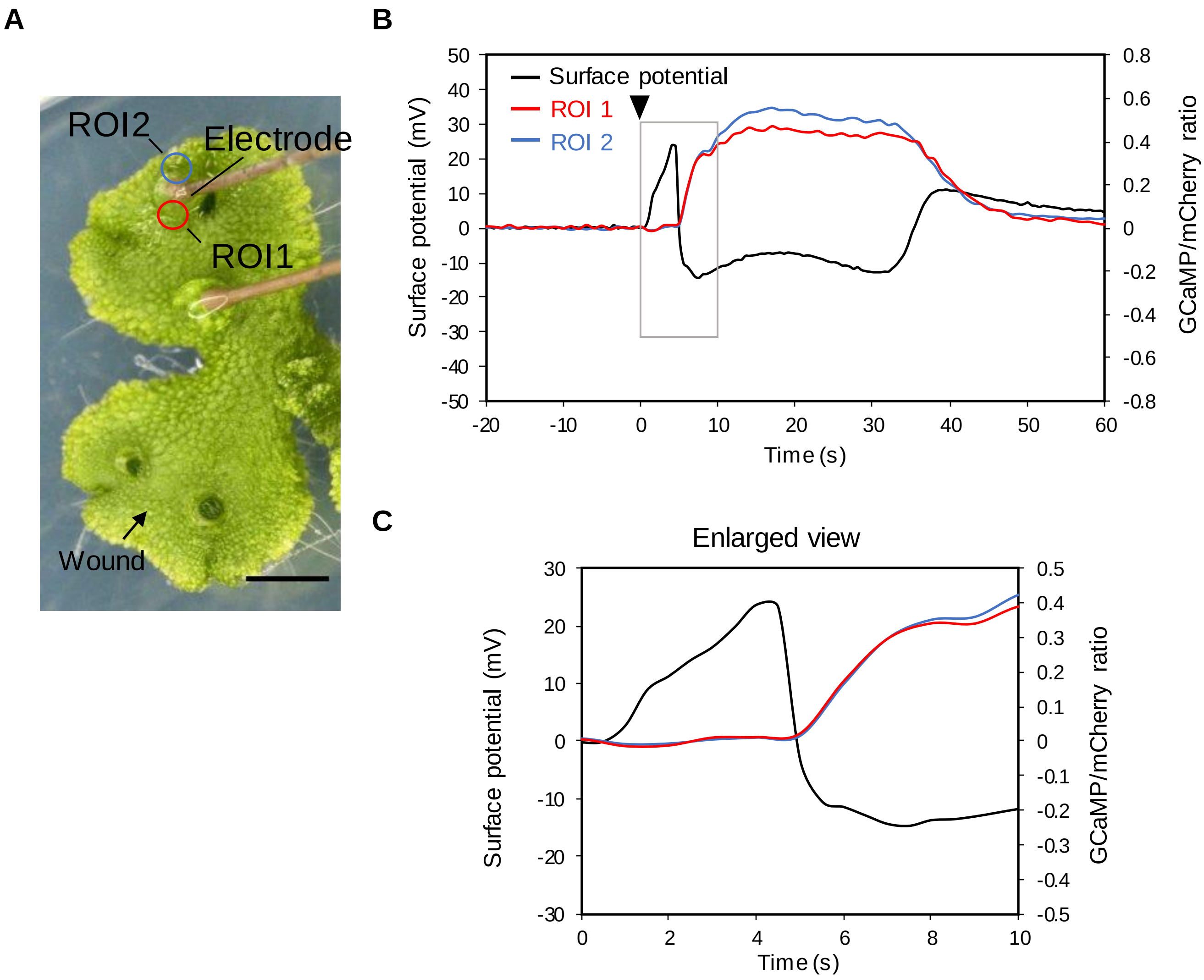


**Figure S6. Simultaneous measurements of $[Ca^{2+}]_{cyt}$ and surface potential.** (A) Location of electrodes and ROI1 and ROI2 where GCaMP6f fluorescence intensity was measured. Scale Bar = 3 mm. (B) Recording of fluorescence intensity of GCaMP6f at ROI1, ROI2 and surface potential at electrode. The triangular arrow indicates the time of wounding. (C) An enlarged view of the graph shown in Figure B, focusing on the time interval from 0 to 10 s.

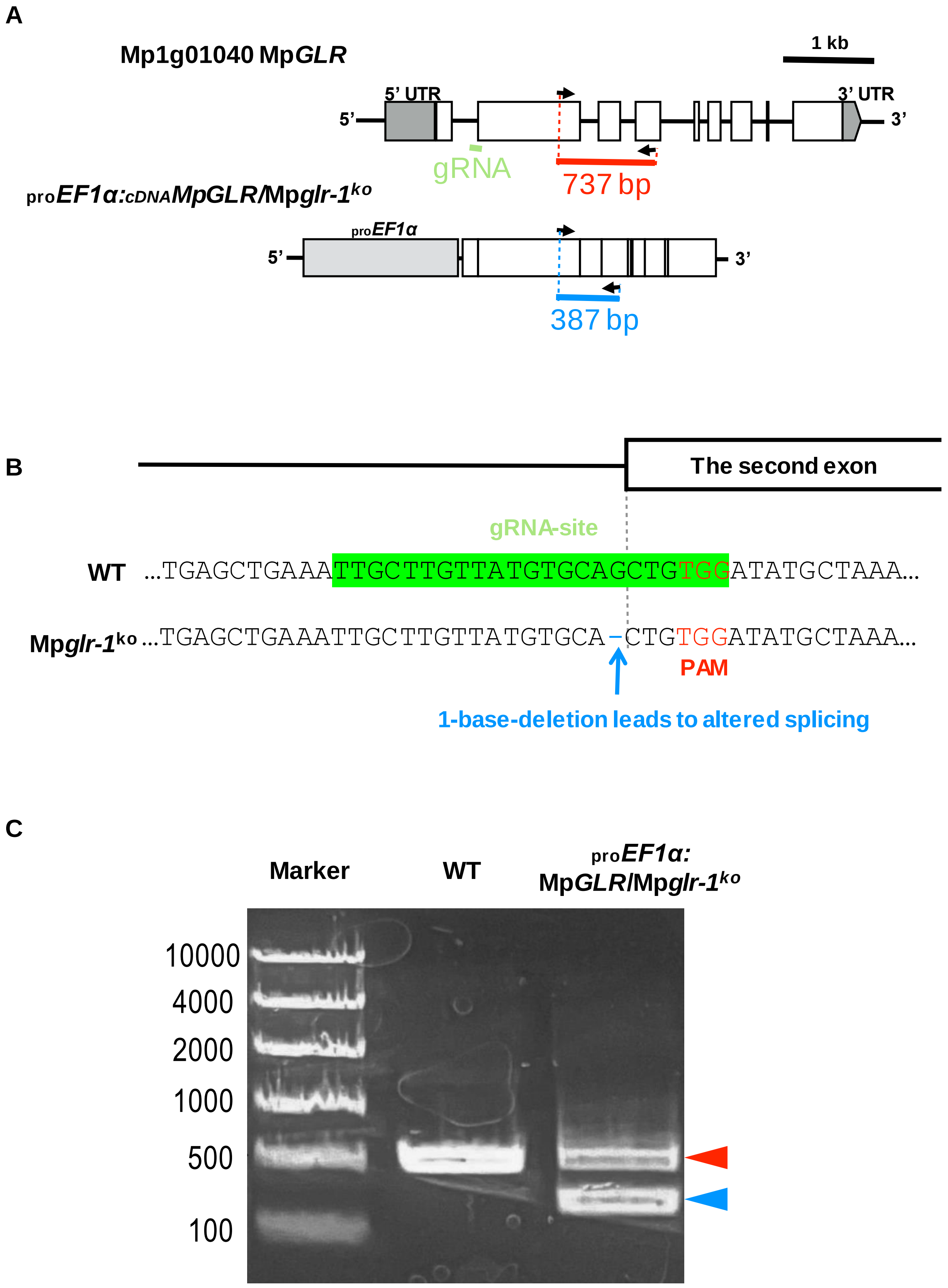


**Figure S7. Summary of Mp*glr*$^{ko}$ mutant and the complementation line**. (A) Schematic diagram of Mp*GLR* gDNA and cDNA and primer positions used in PCR. (B) Genome editing position of Mp*glr-1*$^{ko}$. (C) Electrophoresis of PCR products in the wild-type (WT) and the *proEF1α*:Mp*GLR*/Mp*glr-1*$^{ko}$ complementation line. Red and blue arrows indicate PCR products amplified from Mp*GLR* gDNA and cDNA, respectively.
